# Robustness of respiratory rhythm generation across dynamic regimes

**DOI:** 10.1101/549444

**Authors:** Jonathan E. Rubin, Jeffrey C. Smith

## Abstract

A central issue in the study of the neural generation of respiratory rhythms is the role of the intrinsic pacemaking capabilities that some respiratory neurons exhibit. The debate on this issue has occurred in parallel to investigations of interactions among respiratory network neurons and how these contribute to respiratory behavior. In this computational study, we demonstrate how these two issues are inextricably linked. We use simulations and dynamical systems analysis to show that once a conditional respiratory pacemaker, which can be tuned across oscillatory and non-oscillatory dynamic regimes in isolation, is embedded into a respiratory network, its dynamics become masked: the network exhibits similar dynamic properties regardless of the conditional pacemaker node’s tuning, and that node’s outputs are dominated by network influences. Furthermore, the outputs of the respiratory central pattern generator as a whole are invariant to these changes of dynamical properties, which ensures flexible and robust performance over a wide dynamic range.

**Author summary:** Breathing movements in mammals are generated by brainstem respiratory central pattern generator (CPG) networks, which incorporate an excitatory oscillator located in the pre-Bötzinger Complex (preBötC) that can exhibit autorhythmic behavior. To understand how these autorhythmic properties impact CPG network dynamical performance, we performed computational studies with an established modeling framework to systematically analyze network behavior when the preBötC excitatory neurons’ intrinsic dynamics are tuned to operate in autorhythmic versus non-autorhythmic regimes. Both of these regimes enable rhythmic activity of the CPG network, and we show that the rhythm and its responses to various manipulations are preserved across the tunings of intrinsic properties of the preBötC component. Correspondingly, the emergence of behaviorally appropriate rhythmic patterns of network activity is maintained across preBötC regimes, accompanied by an expansion of the ranges of network output frequencies and amplitudes beyond those attainable with either preBötC regime alone. These results lead to the novel conclusion and concept that the dynamical operation of the CPG is functionally highly robust with respect to the rhythmogenic state of the preBötC excitatory circuits, which could represent a key property for preserved respiratory function across varying conditions and demands on network performance.

## 1 Introduction

A variety of neuronal circuits, including a range of brainstem and spinal cord central pattern generators (CPGs) in many species, exhibit rhythmic activity patterns. In many CPGs, these patterns consist of sequential activations of different neuronal populations that interact through synaptic connections [1, 2]. Significant effort has gone into exploring, using experimental and theoretical methods, the extent to which the intrinsic bursting or pacemaking capabilities of neurons within these populations are responsible for the existence of the network rhythms in which they participate. For example, experimental studies have established the existence of intrinsically bursting neurons in the respiratory pre-Bötzinger complex (preBötC), the inspiratory oscillator within the mammalian brainstem [3], and certain experimental manipulations of burst-supporting conductances in these neurons have eliminated respiratory rhythms [4-7].

Although these and a wide range of other investigations have explored burst generation mechanisms in respiratory preBötC neurons and have debated the role of this burst generation capability in respiratory rhythms across a variety of conditions [8, 9], these studies have not answered a key question: What happens to this intrinsic bursting when the burst-capable neurons are embedded within the full network with which they interact synaptically? In fact, it remains unknown whether the intrinsic bursting capabilities of subsets of preBötC neurons affect the dynamics that these neurons actually exhibit once they are embedded within a synaptically interconnected respiratory circuit and how this bursting capability contributes to the properties of the circuit’s rhythmic outputs.

In this study, we use highly reduced models of the respiratory CPG, composed of a small number of units representing important neuron populations [10], to highlight some key ideas relating to these issues. We model the burst-capable neurons in the network using differential equations that incorporate a representation of the persistent sodium current, which supports an established oscillatory burst generating mechanism and has been well documented in these neurons experimentally [6, 11-13]. Basic principles of dynamical systems theory imply, however, that any alternative mechanisms providing a similar voltage-dependence of activity and autorhythmic capability would yield similar results [14], and hence the ideas that we illustrate are not dependent on a specific bursting mechanism. In particular, our key finding is that network rhythmicity extends smoothly and robustly across different dynamic regimes of the intrinsically burst-capable neurons in the network. Within the multi-phase rhythmic outputs that emerge, these neurons’ outputs can become dominated by network influences. While network rhythms exhibit similar features whether the tuning of these neurons renders them intrinsically burst-capable or not, the maintenance of output function across different tunings of intrinsic dynamics yields a broadened dynamic range of network output properties.

## 2 Results

### 2.1 Properties of pre-I neuronal unit dynamics are masked within network rhythms

We simulated a highly reduced rhythm-generating network model consisting of ordinary differential equations representing coupled neuronal units. We based our network connectivity on published models for the generation of rhythmic activity patterns in respiratory circuits [5] and we tuned the network to produce rhythmic outputs qualitatively similar to the experimentally observed tri-phasic rhythmic respiratory pattern, including the inspiratory (I), the post-I expiratory, and the subsequent augmenting-expiratory (aug-E) stages. The circuit depicted (Fig. 1) consists of a preBötC excitatory (glutamatergic) unit (often referred to as pre-I/I because its firing occurs before and during the I phase, but abbreviated as pre-I throughout this work) representing rhythmogenic neurons in the preBötC, a brainstem region that can exhibit intrinsic oscillatory dynamics and drives I activity, along with inhibitory units named after the timing within the respiratory cycle at which their activity occurs (early-I, post-I and aug-E). Among these, it is thought that critical subpopulations of post-I and aug-E inhibitory neurons in respiratory CPG circuits reside in the BötC region adjacent to the preBötC [5, 8], and these inhibitory neurons have been shown to provide strong inhibition to the preBötC to regulate inspiratory rhythm [15]. In some simulations, as noted in the corresponding text, we also included a unit representing the recently identified glutamatergic post-inhibitory complex (PiCo) [16]. Within this framework, we tuned the parameters in the equations for each unit to represent a distinct activity profile experimentally identified to arise in a corresponding subpopulation of respiratory neurons during the respiratory cycle (see Fig. 4C,D) [5]. In each of these populations, neurons become active and fall silent in a synchronized way but need not spike in synchrony, and hence we omitted spiking currents in our model units.

**Fig 1.**
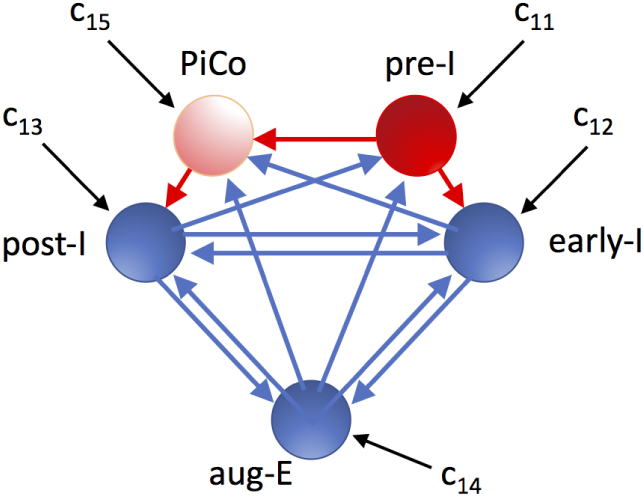
Schematic illustration of the respiratory network configuration considered. Red is used for excitatory units and their outputs, while blue denotes inhibitory units and their outputs. Black arrows correspond to tonic drives to units in the network. The PiCo is only included in a subset of our simulations, as indicated in the text.

**Fig 4.**
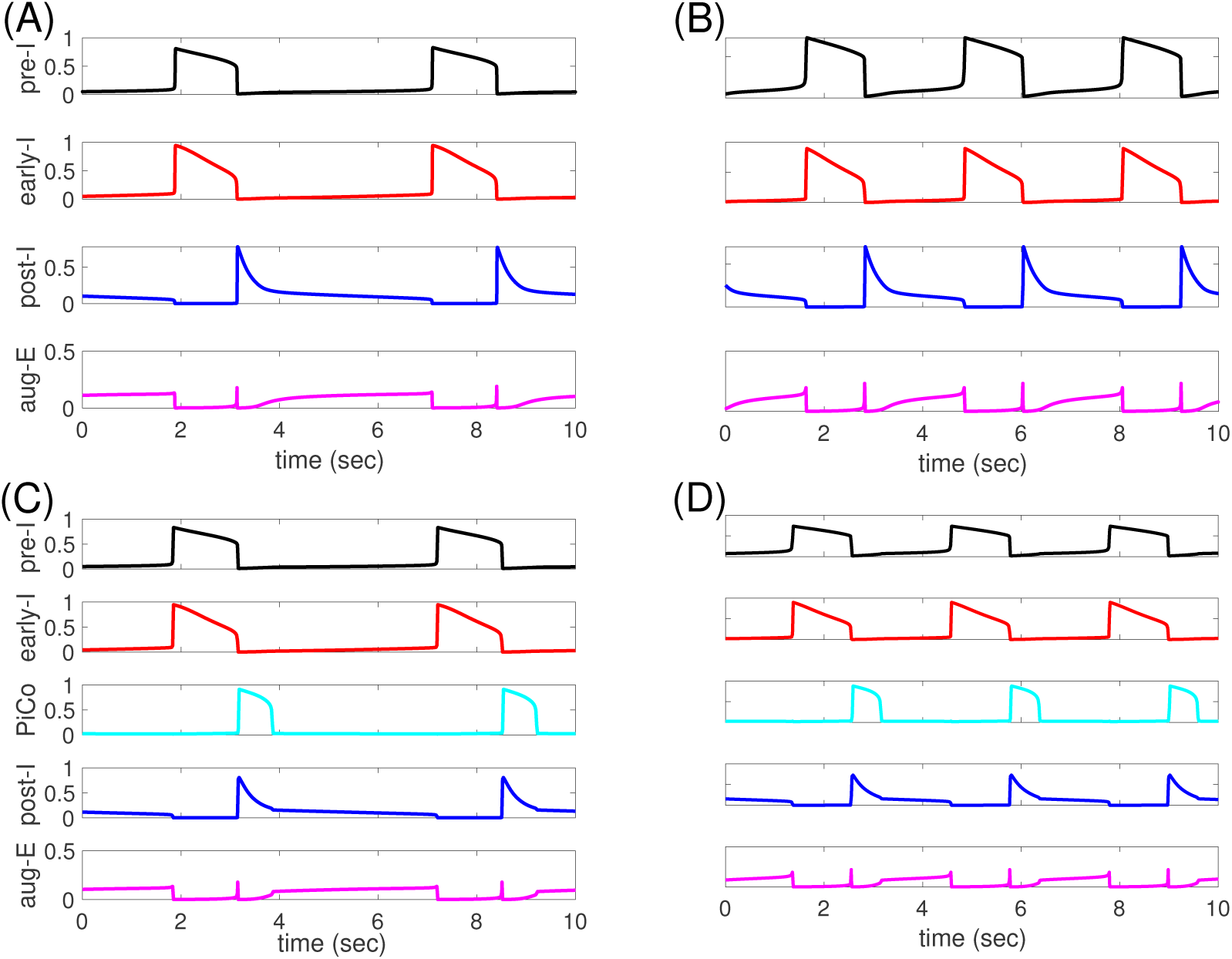
Basic model behavior. (A-B): Time courses of outputs of all 4 neurons (with PiCo silent and omitted) in the network rhythm when (A) pre-I is intrinsically oscillatory (*c*_11_ = –0.03) or (B) pre-I is intrinsically tonic (*c*_11_ = 0.01). Black: pre-I. Red: early-I. Blue: post-I. Magenta: aug-E. (C-D): Time courses of outputs of all 5 neurons in the network rhythm, including PiCo (*c*_15_ = 0.045), when (C) pre-I is intrinsically oscillatory (*c*_11_ =–0.03) or (D) pre-I is intrinsically tonic (*c*_11_ = 0.01). Black: pre-I. Red: early-I. Cyan: PiCo. Blue: post-I. Magenta: aug-E.

Within the respiratory rhythm-generating circuit, some members of the pre-I population in the preBötC have been found to be capable of producing rhythmic bursting activity even when isolated from synaptic inputs, at least over some range of constant input currents or extracellular potassium concentrations [3, 17, 18]. We simulate the conditional bursting property of pre-I neurons by endowing the pre-I unit’s voltage equation with a persistent sodium current, *I*_*NaP*_, which is an established feature of preBötC excitatory neurons [6, 11, 13] and provides a voltage-controlled relaxation-oscillator type of rhythmic oscillation. For relatively low values of the pre-I unit’s tonic drive parameter, *c*_11_, the unit’s voltage-nullcline is cubic and features a fixed point on its hyperpolarized (or left) branch, corresponding to a quiescent state. As *c*_11_ is increased, the fixed point crosses to the middle branch and a bifurcation occurs (an Andronov-Hopf, or A-H, bifurcation with a canard explosion [19], after which the unit will, in isolation from synaptic inputs from other units, produce relaxation oscillations. Finally, with additional increases in *c*_11_, the nullcline switches from cubic to monotonically increasing and another (A-H) bifurcation occurs, producing the loss of oscillations and the appearance of a depolarized fixed point, corresponding to a tonically activated state (Fig. 2). Note that the actual numerical values of *c*_11_ used here are given in arbitrary units.

**Fig 2.**
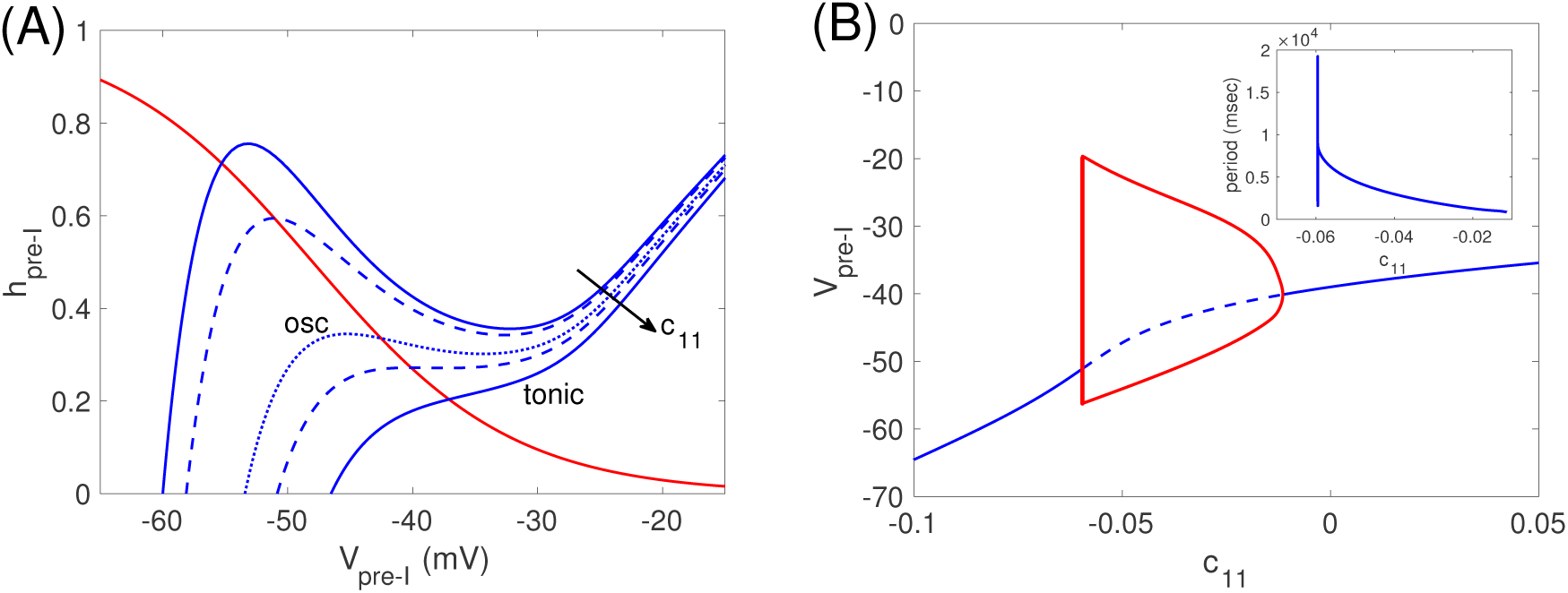
Pre-I tuning. (A): The nullcline structure for the pre-I unit in the preBötC depends on its tonic drive parameter, *c*_11_. Blue curves denote the *V*_*pre–I*_-nullcline for several values of *c*_11_, with different line patterns used for contrast enhancement. The red curve is the *h*_*pre–I*_-nullcline, which is independent of *c*_11_. (B): Bifurcation diagram for the pre-I unit with respect to *c*_11_. Solid (dashed) curve denotes stable (unstable) equilibria. Red curves are max and min voltages along a family of stable periodic orbits; these grow extremely rapidly in amplitude when they first appear, near *c*_11_ = *−*0.060. Pre-I is oscillatory where these orbits exist, on *c*_11_ ∈ (*−*0.060, *−*0.011). Inset shows dependence of periods on *c*_11_; period jumps sharply from 0 at the onset of oscillations, drops abruptly, and then decays more gradually as *c*_11_ increases.

Once this pre-I unit is embedded within the full respiratory network (as shown in Fig. 1, but without the PiCo), its intrinsic properties are masked, or superseded by the network dynamics. There are multiple manifestations of this form of masking. First, at the low end of the *c*_11_ range over which the isolated pre-I unit generates oscillations, the coupled network does not produce oscillatory dynamics but rather settles into a tonic activity regime in which the post-I unit maintains a steady elevated activation level while the pre-I and early-I units exhibit low activation levels. Second, the ranges of periods and amplitudes that the pre-I unit exhibits during its intrinsic oscillations differ from those that it produces when operating within the oscillatory dynamics of the full network (Fig. 3). The sensitivity of the pre-I oscillation period and amplitude to changes in *c*_11_ is also altered by the embedding.

**Fig 3.**
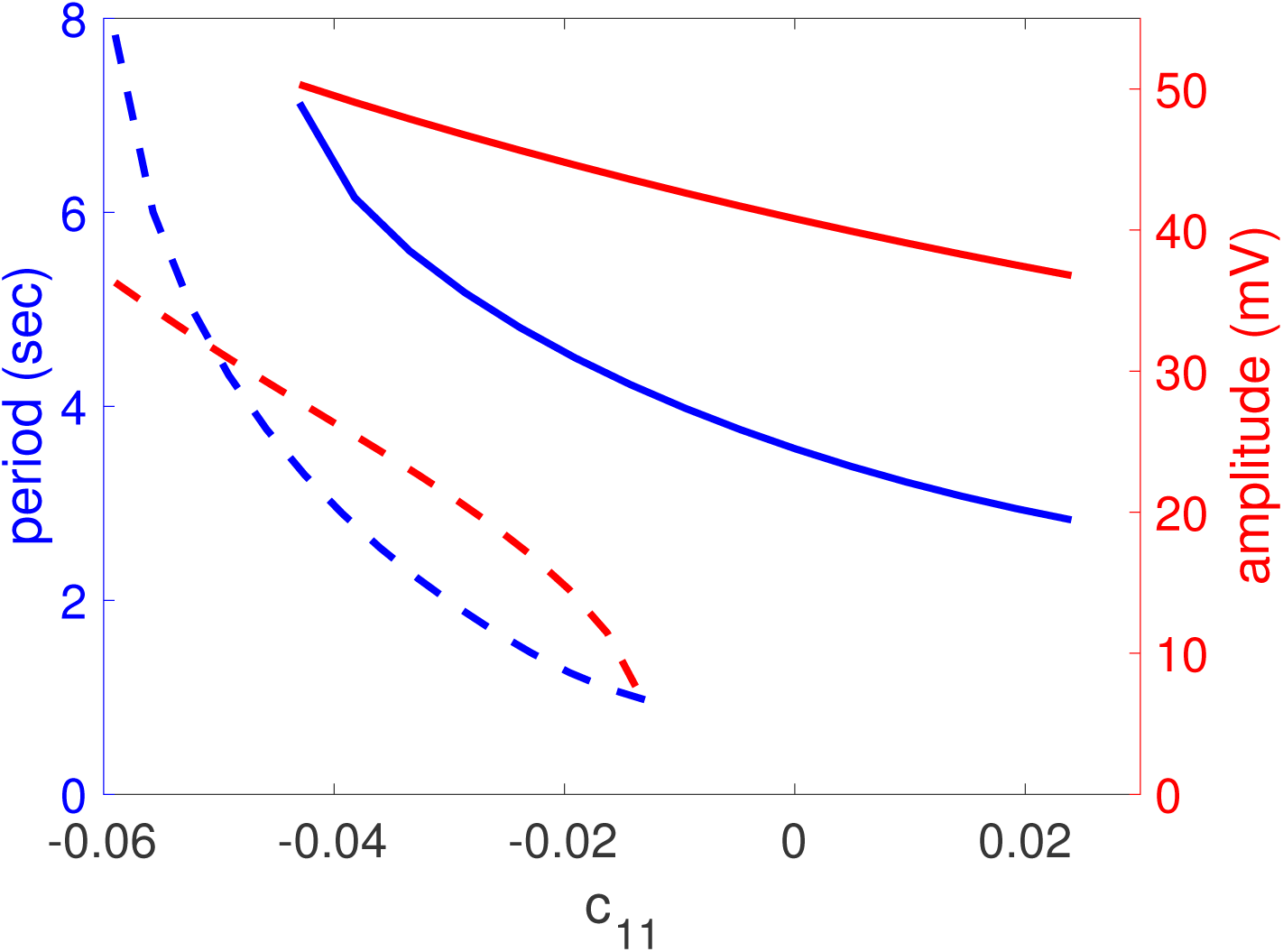
Pre-I output properties are altered by embedding within the full respiratory network. Dashed curves show the period (blue) and amplitude (red) of oscillations exhibited by the pre-I unit in isolation, over the range of *c*_11_ values for which it intrinsically generates oscillatory behavior. Solid curves illustrate the same quantities for the pre-I unit when it is embedded in the full model respiratory network, over the range of *c*_11_ values for which the network oscillates.

Interestingly, the activity pattern of the synaptically connected respiratory network remains remarkably similar regardless of whether *c*_11_ is tuned to yield oscillatory or tonic pre-I dynamics in isolation. These network activity patterns are shown in Figure 4A,B for representative *c*_11_ values, one per regime of intrinsic pre-I dynamics, in terms of nondimensional variables (scaled between 0 and 1, with the same scaling used across all *c*_11_) representing the output of the pre-I, early-I, post-I, and aug-E units that comprise the network, when PiCo is silent. In both cases, pre-I and to a lesser extent early-I output slowly ramps up before pre-I and early-I activity levels surge together, corresponding to the onset of inspiration. Pre-I and early-I then gradually decline before an abrupt take-over by post-I unit activity and corresponding inhibition of the two I units, marking the phase switch to expiration. After this switch, post-I output declines, disinhibiting aug-E, the output of which ramps up, yielding what is known as the late-expiratory or E-2 phase, with aug-E output exceeding and inhibiting but not completely suppressing post-I. This similarity in the pattern of network dynamics across different pre-I unit tunings represents another form of masking, since the intrinsic tuning of the pre-I unit dynamics becomes hidden and cannot be deduced from observation of the network activity pattern.

A comparison between the examples shown reveals that the periods do differ somewhat. We will next explore frequency responses to parameter changes across the two regimes of pre-I intrinsic dynamics, and later we will consider network dynamics when PiCo activity is included (Fig. 4C,D).

### 2.2 Period and amplitude responses to reductions of *g*_*NaP*_

In the model that we are considering, we use a persistent sodium current, *I*_*NaP*_, to allow the possibility of intrinsic rhythmicity in our pre-I unit. In experiments, addition of the *I*_*NaP*_ blocker riluzole to the perfusate of an *in situ* rat preparation that preserved the intact respiratory network yielded little change in respiratory period but caused the output of the phrenic nerve, which corresponds to the excitatory output from the inspiratory neurons in the respiratory circuit, to decrease in amplitude [5]. This decrease occurred roughly linearly with riluzole concentration, reaching about 1/3 of the original amplitude with the maximal riluzole dose applied.

We tested the dependence of the period of our network rhythm (*T*), the duration of its inspiratory (*T*_*I*_) and expiratory (*T*_*E*_) phases, and the amplitude of the pre-I unit’s output on the strength of *I*_*NaP*_, represented by its maximal conductance *g*_*NaP*_. Moreover, we evaluated this dependence over a range of values of the pre-I tonic drive parameter *c*_11_ spanning across regimes of both oscillatory and tonic pre-I intrinsic dynamics. Note that the value of *c*_11_ at which the pre-I unit’s intrinsic dynamics changes from oscillatory to tonic depends on *g*_*NaP*_. Since this transition occurs at an A-H bifurcation as shown in Fig. 2B, we used the continuation capabilities of XPPAUT [20] to follow this bifurcation in the two parameters (*g*_*NaP*_, *c*_11_), thereby generating a transition curve.

We found that the quantitative properties of the network rhythm did depend on both *g*_*NaP*_ and *c*_11_ (Fig. 5). For example, *T* was sensitive to *g*_*NaP*_ over a range of *c*_11_ values but less so for sufficiently large *c*_11_. A sufficient reduction of *g*_*NaP*_ prevented network output of a rhythmic pattern for most of the range of *c*_11_ considered, although for *c*_11_ sufficiently large, rhythmic outputs persisted down to *g*_*NaP*_ = 0 *nS*, despite the lack of intrinsic pre-I rhythmicity at this *c*_11_ level. Wider ranges of *T, T*_*I*_, and *T*_*E*_ values could occur for relatively small (i.e., relatively negative) *c*_11_ than for large *c*_11_ (Fig. 5A,C,D), while amplitude changed more with *g*_*NaP*_ when *c*_11_ was large (i.e., near 0 or positive, Fig. 5B). Importantly, however, when *g*_*Na*_ and *c*_11_ were varied from one side of the oscillatory-tonic transition curve (red curves in Fig. 5) to the other, the transition in pre-I intrinsic dynamics caused absolutely no associated jump or shift in the quantitative properties of the network rhythm (Fig. 5).

**Fig 5.**
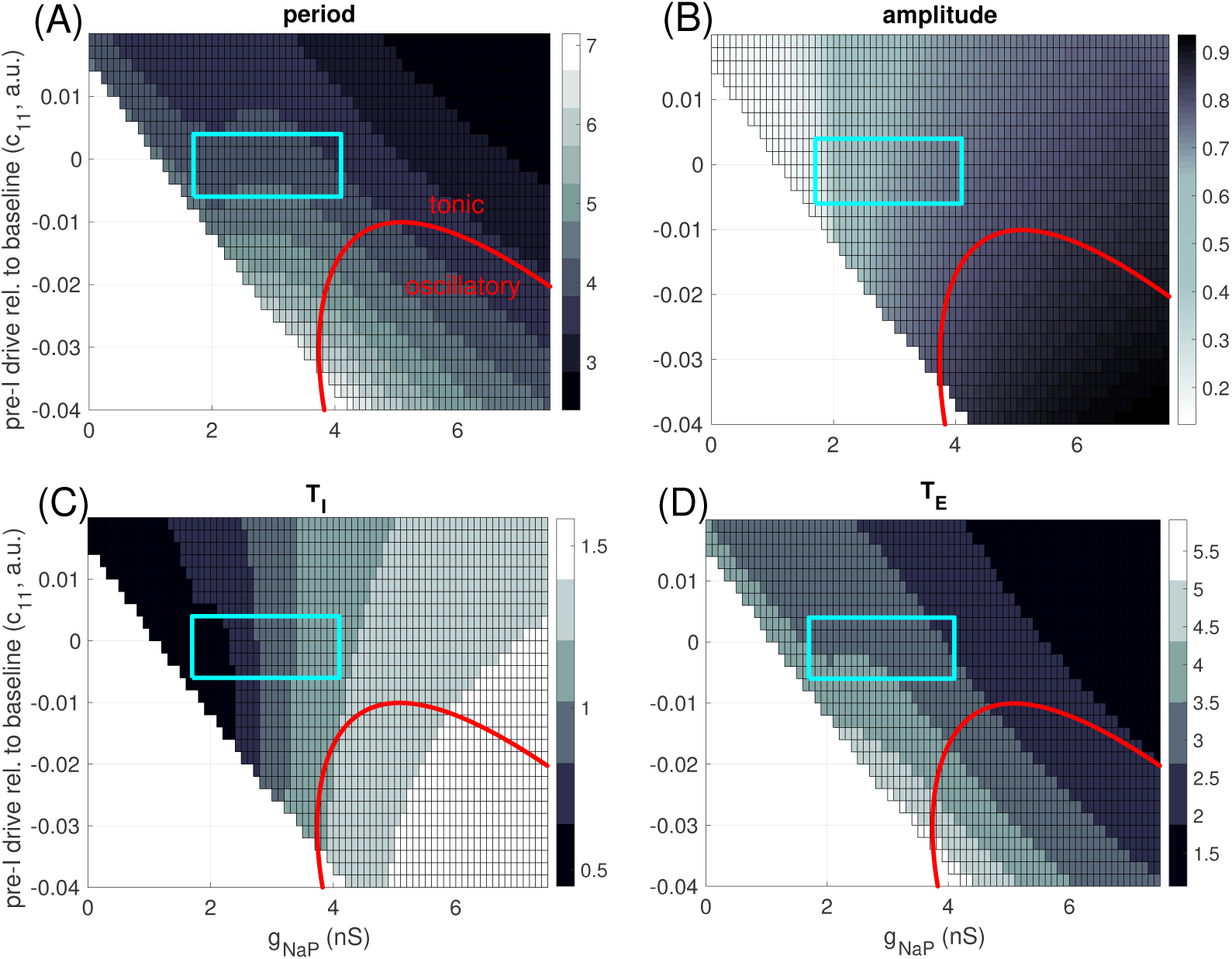
Parameter-dependence of model output. (A) Grey scale in boxes codes period of full rhythm, with longest period given by palest shades (color bar, periods in seconds). For parameter values in the unboxed white region, the network is not rhythmic. For parameter values below the red curve, the pre-I neuron is intrinsically oscillatory. (B) Grey scale codes amplitudes, with largest amplitude given by darkest shades (color bar, amplitudes measured as maximum minus minimum of pre-I output, *f* (*V*_1_). Red curve as in upper left. (C) Grey-coded inspiratory phase duration (seconds), measured as time between increased and subsequent decreases of *V*_1_ through a threshold of -35 mV. (D) Grey-coded expiratory phase duration (seconds), measured as the period minus the inspiratory phase duration.

Finally, we compared the dependence of the model on *g*_*NaP*_ to that found experimentally [5]. We found that over a range of *c*_11_ values straddling 0 (*c*_11_ ∈ [*−*0.006, 0.004]), we could select a range of *g*_*NaP*_ values (*g*_*NaP*_ ∈ [1.7 *nS,* 4.1 *nS*]) such that when *g*_*NaP*_ was reduced through this range, the model provided an excellent agreement with the experimentally observed [5] invariance of period and 2/3 drop in amplitude of inspiratory output (blue boxes, Fig. 5A,B; amplitude range approximately 0.23 to 0.75). In fact, there was some decrease in *T*_*I*_ over this range (Fig. 5C), but it was compensated by an increase in *T*_*E*_ (Fig. 5D) to yield little net change in *T* (Fig. 5A).

### 2.3 Dynamic mechanisms of network rhythms do not depend on intrinsic tuning of preBötC excitatory neurons

Although the network’s rhythmic activity pattern is similar for both oscillatory and tonic intrinsic preBötC unit dynamics, it remains possible that this intrinsic tuning could affect the detailed dynamic mechanisms by which this pattern is produced, potentially leading to different dynamics in response to certain inputs or perturbations. Since we use model units with dynamics that can be visualized in the phase plane to model each population in the network, we can represent the network oscillation mechanisms geometrically and compare them between the two cases. To do so, although the full network model consists of an eight-dimensional dynamical system, we consider the network outputs projected to various two-dimensional phase planes, each defined using the two variables for one specific unit (Figs. 6, 7). Within these phase planes, we plot the nullclines, or curves of zero rate of change, for each of these two variables. Since each unit’s voltage equation depends on the outputs of the other units that are coupled to it synaptically, the location of each projected voltage nullcline varies with the activity of each presynaptic unit. We plot the nullclines at several key moments within a network activity cycle that are particularly helpful for understanding the network dynamics. We also plot the projection of the network output trajectory to each phase plane. Because voltage changes much more rapidly than the persistent sodium inactivation variable, these trajectory projections generally lie on appropriate voltage nullclines except during brief excursions when the network output rapidly changes, which occur in the transitions between inspiratory and expiratory phases. Note that we omit the early-I unit from this discussion, because it always settles to a stable fixed point corresponding to the level of input that it receives, and it lacks the auto-rhythmic capability needed to initiate a transition to the I phase.

**Fig 6.**
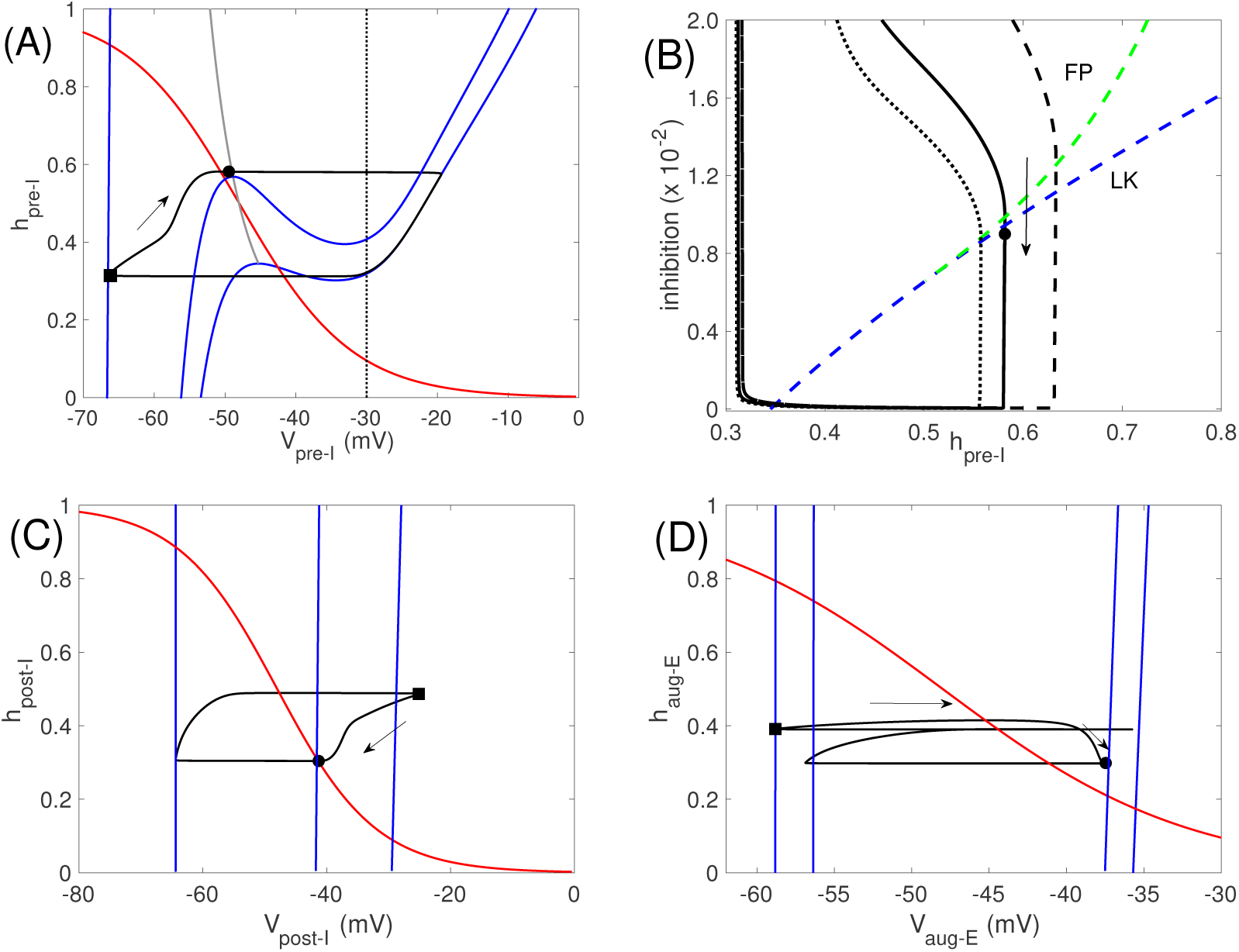
Network oscillation mechanisms when pre-I is intrinsically oscillatory, with *c*_11_ = *−*0.03. (A) Phase plane (*h*_*pre–I*_ versus *V*_*pre–I*_) for the pre-I unit. Black curve is the model output projected to this plane. Solid blue curves are *V*_*pre–I*_-nullclines. The lower right nullcline corresponds to 0 inhibition to pre-I, the upper left nullcline (which continues out of the image) to the maximal inhibition received by pre-I (0.0965), and the middle nullcline to the inhibition level received by pre-I (0.09) when it begins its escape to the active phase by hitting the curve of knees (grey). The red curve is the *h*_*pre–I*_-nullcline, which is independent of inhibition level. The dotted black curve denotes the *V*_*pre–I*_ value at which the output of pre-I is half of its maximum value. Solid black circles and squares correspond to time points when inhibition to the pre-I unit is 0.09 and 0.0965, respectively. (B) Same solution projected to the plane in which the inhibition to pre-I is plotted versus *h*_*pre–I*_ (black solid). As inhibition wears off, the trajectory first crosses the curve of pre-I fixed points (FP, green dashed) and then crosses the curve of *V*_*pre–I*_-nullcline left knees (LK, blue dashed), allowing escape (black circle) and initiation of inspiration. Trajectories for reduced/increased inhibition from post-I to pre-I (*b*_31_ = 0.105*/*0.175) are also shown (dotted/dashed black). (C) Phase plane (*h*_*post-I*_ versus *V*_*post-I*_) for the post-I unit. Curves are analogous to those in (A). The *V*_*post-I*_-nullclines shown correspond to minimal inhibition to post-I (far right, inhibition is 0), occurring when the maximal value of *V*_*post-I*_ is achieved along the trajectory (black square, same time point as in (A)); to maximal inhibition to post-I (far left, inhibition is 0.54); and to the inhibition level when pre-I initiates its escape (middle, inhibition is 0.0525, also marked by solid black circle at same time point as in (A)). (D) Phase plane (*h*_*aug–E*_ versus *V*_*aug–E*_) for the aug-E unit. Curves are analogous to those in (A). From right to left, the *V*_*aug–E*_-nullclines shown correspond to minimal inhibition to aug-E (inhibition is 0.05); to inhibition to aug-E at the moment when pre-I initiates its escape (inhibition is 0.0542, also marked by black circle at same time point as in other plots); to inhibition to aug-E when *V*_*pre–I*_ is at its peak (0.283); and to inhibition to aug-E when *V*_*post-I*_ is at its peak (0.35, marked by black square at same time point as in other plots).

The phase plane for the pre-I unit shows that even if the preBötC unit is tuned to be intrinsically oscillatory, at the moment when it receives its maximal level of inhibition, which occurs at the onset of post-I activity (black square in Fig. 6A), its voltage- or *V*_*pre–I*_-nullcline (leftmost blue curve, Fig. 6A) intersects the *h*_*pre–I*_-nullcline in a stable fixed point at large *h*_*pre–I*_, such that the unit cannot activate through *I*_*NaP*_ deinactivation alone. At that moment, *V*_*post-I*_ is at its maximum (black square in Fig. 6C), while *V*_*aug–E*_ is hyperpolarized due to the strong inhibition that the aug-E unit receives from post-I (black square in Fig. 6D). As can be seen in Fig. 6C,D, during the ensuing post-I phase, *V*_*post-I*_ gradually decreases, which occurs due to an inhibitory input from the aug-E unit (which is already present – although very weak – near the onset of the expiratory phase in our model tuning, as can be seen in Fig. 4), and *V*_*aug–E*_ gradually increases; together, these changes form a self-reinforcing feedback loop. As a result, the inhibitory output from the post-I unit drops, the *V*_*pre–I*_ - nullcline position changes, and the value of *V*_*pre–I*_ increases correspondingly (Fig. 6A, arrow).

A key moment in the respiratory cycle is the transition from expiration to inspiration. From the phase plane in Fig. 2, recall that the *V*_*pre–I*_ nullcline has a cubic shape over a range of parameter values when the pre-I unit does not receive any inhibition. Because the inhibition from post-I to pre-I is stronger than the inhibiton from aug-E to pre-I, it follows that once *V*_*post-I*_ is sufficiently reduced, the *V*_*pre–I*_-nullcline becomes cubic. In a phase plane with a cubic voltage nullcline, a unit transitions from the hyperpolarized phase to the depolarized phase when its voltage rises above the left fold or knee of its voltage nullcline. The grey curve in Fig. 6A is the curve of knees for this unit. The *V*_*pre–I*_-nullcline is shown at the moment when *V*_*pre–I*_ reaches this curve of knees (black circle), allowing the transition to inspiration to commence (see also black circles and corresponding voltage nullclines in Fig. 6C,D).

Another view of this transition is provided by plotting the total level of inhibition to the pre-I unit versus *h*_*pre–I*_ (Fig. 6B). The (*V*_*pre–I*_, *h*_*pre–I*_) coordinates of both the knee itself (dashed blue curve) and the fixed point (dashed green curve), where the *V*_*pre–I*_ and *h*_*pre–I*_ nullclines intersect, depend on this inhibition level. The projection of the trajectory would tend to the curve of fixed points if inhibition were held fixed, preventing the onset of inspiration. But the gradual decrease of total inhibition to pre-I carries the trajectory across the curve of knees, allowing inspiration to begin. The pre-I unit jumps to and follows the right branch of its uninhibited voltage nullcline, corresponding to inspiration (rightmost blue curve, Fig. 6A), while the post-I and aug-E units become strongly inhibited and jump to corresponding inhibited voltage nullclines (Fig. 6C,D). Finally, once *V*_*pre–I*_ becomes close enough to its synaptic threshold (black dotted line, Fig. 6A), the inhibition to post-I and aug-E weakens and eventually both of their nullclines shift to depolarized voltages. Both units attempt to activate but post-I wins, yielding a transient spike and subsequent hyperpolarization of the aug-E unit and the transition to expiration, completing the cycle.

Note that the only possible moment in this process when the intrinsic oscillation capability of the pre-I unit could have mattered was at the onset of inspiration, which depends on the cubic shape of the *V*_*pre–I*_-nullcline. Yet examining similar phase plane representations when the pre-I unit is tuned to be intrinsically tonic shows that even in this tuning, the *V*_*pre–I*_-nullcline is still cubic at the moment of onset of inspiration and the mechanism underlying the transition to inspiration is the same (Fig. 7A,B). That is, even though this nullcline becomes monotonic in this tuning in the absence of inhibition, some inhibition is still present when the network transitions to inspiration, which causes the *V*_*pre–I*_-nullcline to be cubic. This mechanism is robust; the *V*_*pre–I*_-nullcline has a cubic shape, with a corresponding left knee, over a wide range of inhibition levels to the pre-I unit (dashed blue curve, Fig. 7B). Not surprisingly, the phase plane projections for the post-I and aug-E units are qualitatively identical between the two pre-I unit tunings as well (data not shown).

**Fig 7.**
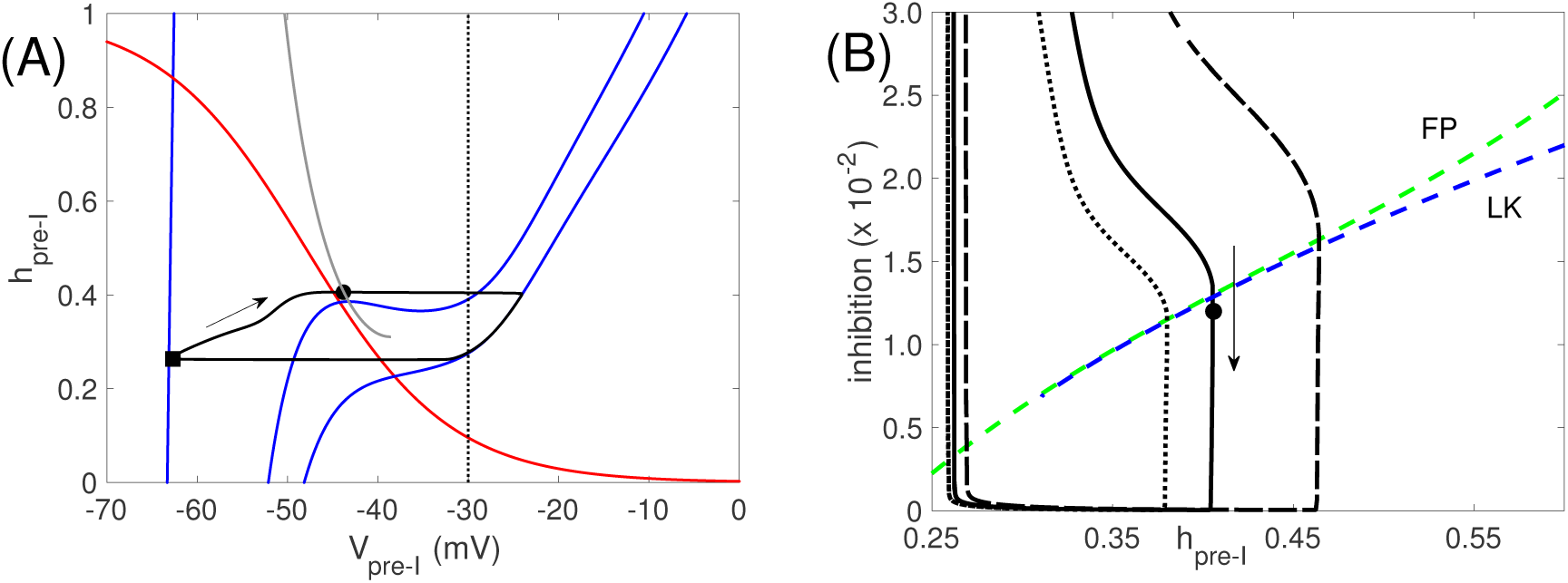
Network oscillation mechanisms when pre-I is intrinsically tonic, with *c*_11_ = 0.01. Mechanisms are identical to the case when pre-I is intrinsically oscillatory. (A) Analogous curves to Fig. 6A. Maximal inhibition is 0.088 (black square), minimal inhibition is 0, inhibition at escape is 0.012 (black circle). Note that the grey curve of knees terminates when the nullcline becomes monotone (nullcline not shown). (B) Analogous curves to Fig. 6B. The termination of the curve of left knees is also evident here (around inhibition of 0.7 *×*10^−2^).

### 2.4 Parameter dependence of network rhythms does not depend on intrinsic tuning of preBötC excitatory neurons

The fact that the same dynamic mechanisms underlie network rhythmicity across parameter tunings explains the insensitivity of output period, phase durations, and amplitude to the switch in intrinsic preBötC dynamics as *g*_*nap*_ and *c*_11_ are varied (Fig. 5). Similarly, we can expect the changes of the network rhythm properties with variations in other network parameters to be independent of intrinsic preBötC tuning. Although we already explored variations in *c*_11_, we first plotted period (*T*) and inspiratory (*T*_*I*_) and expiratory (*T*_*E*_) phase durations versus *c*_11_ for comparison with effects of other parameter variations (Fig. 8A). As seen previously (Fig. 5), there was no abrupt change as the preBötC tuning switched from intrinsically oscillatory to intrinsically tonic. The period *T* decreased as *c*_11_ increased, via a shortening of *T*_*E*_ as also observed experimentally with preBötC stimulation (e.g., [21]), from a maximum of just over 7 seconds to a mimimum of about 3 seconds.

**Fig 8.**
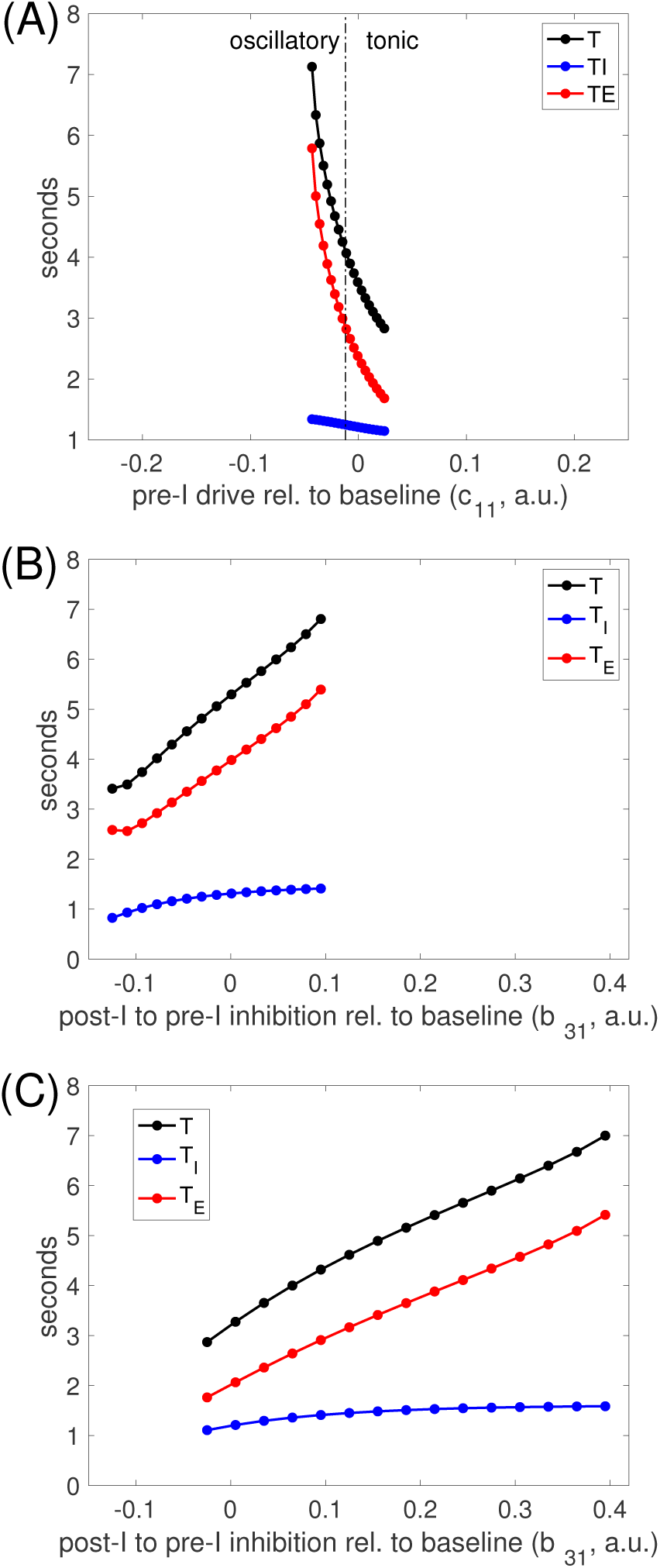
Input to the preBötC pre-I node controls period by modulating *T*_*E*_. (A): Period (black), inspiratory duration (blue), and expiratory duration (red) versus drive to pre-I, *c*_11_, over the range of *c*_11_ for which network rhythms were maintained. Values of *c*_11_ where the pre-I unit is intrinsically oscillatory or tonic are separated (dash-dotted black line) and labeled. (B-C): Same curves versus the value of inhibition from post-I to pre-I, *b*_31_, relative to its baseline value (0.125; here labeled as 0) over the range where network rhythmicity is maintained; results are shown when the pre-I is intrinsically oscillatory (B), with *c*_11_ = –0.03, and tonic (C), with *c*_11_ = 0.01. Increasing *b*_31_ prolongs the time until pre-I activity can rise, leading to longer expiratory phase durations and periods.

Next, since the role of inhibition in respiratory rhythmicity has received significant recent attention [8, 15, 21-24], we considered the effects of varying the strength of the inhibition from the post-I to the pre-I unit, *b*_31_. This parameter variation produced similar effects whether the pre-I unit was tuned to be intrinsically oscillatory (Fig. 8B) or tonic (Fig. 8C): increases in *b*_31_ produced increases in overall period, largely via increases in *T*_*E*_ but with a small accompanying increase in *T*_*I*_, again yielding a range of periods between about 3 and 7 seconds. Although the stronger maximal inhibition allowed *h*_*pre–I*_ to increase faster than with weaker inhibition and therefore allowed the pre-I unit to cross its curve of left knees and activate at a larger inhibition level than previously (dashed black curve, Fig. 6B, Fig. 7B), the amount of time for inhibition to decay from its larger maximum value until it hit the curve of left knees was still longer than the escape time with weaker inhibition, leading to an overall longer *T*_*E*_. The larger value of *h*_*pre–I*_ at the onset of inspiration in this case yielded the small increase in *T*_*I*_. Similarly, reducing *b*_31_ allowed an earlier crossing of the pre-I curve of left knees and shorter *T*_*E*_ (dotted black curve, Fig. 6B, Fig. 7B).

After these manipulations of parameters associated with the pre-I unit, we varied the drive levels to the other units in the network, one by one, to analyze their abilities to tune network oscillation frequency. In all three cases, these parameter changes produced qualitatively similar results across both tunings of the pre-I unit (Fig. 9). Increases in the drive to the early-I unit, *c*_12_, caused decreases in network period via decreases in *T*_*E*_. In contrast, increases in *c*_13_ (post-I drive) and *c*_14_ (aug-E drive) both increased *T,* again via changes in *T*_*E*_. Interestingly, the period was much more sensitive to *c*_14_ than to *c*_13_ (also consistent with previous modeling [10]). This sensitivity emerged because in the baseline tuning, the aug-E unit activation was much more modest than the post-I activation, with the aug-E output much farther from maximal. Thus, changes in drive to aug-E influenced the inhibition level to the pre-I unit more than changes in drive to post-I. Finally, because the baseline period was shorter for the intrinsically tonic than the intrinsically oscillatory pre-I tuning, the range of periods in all cases extended to larger values, producing a wider overall range, for the oscillatory regime.

**Fig 9.**
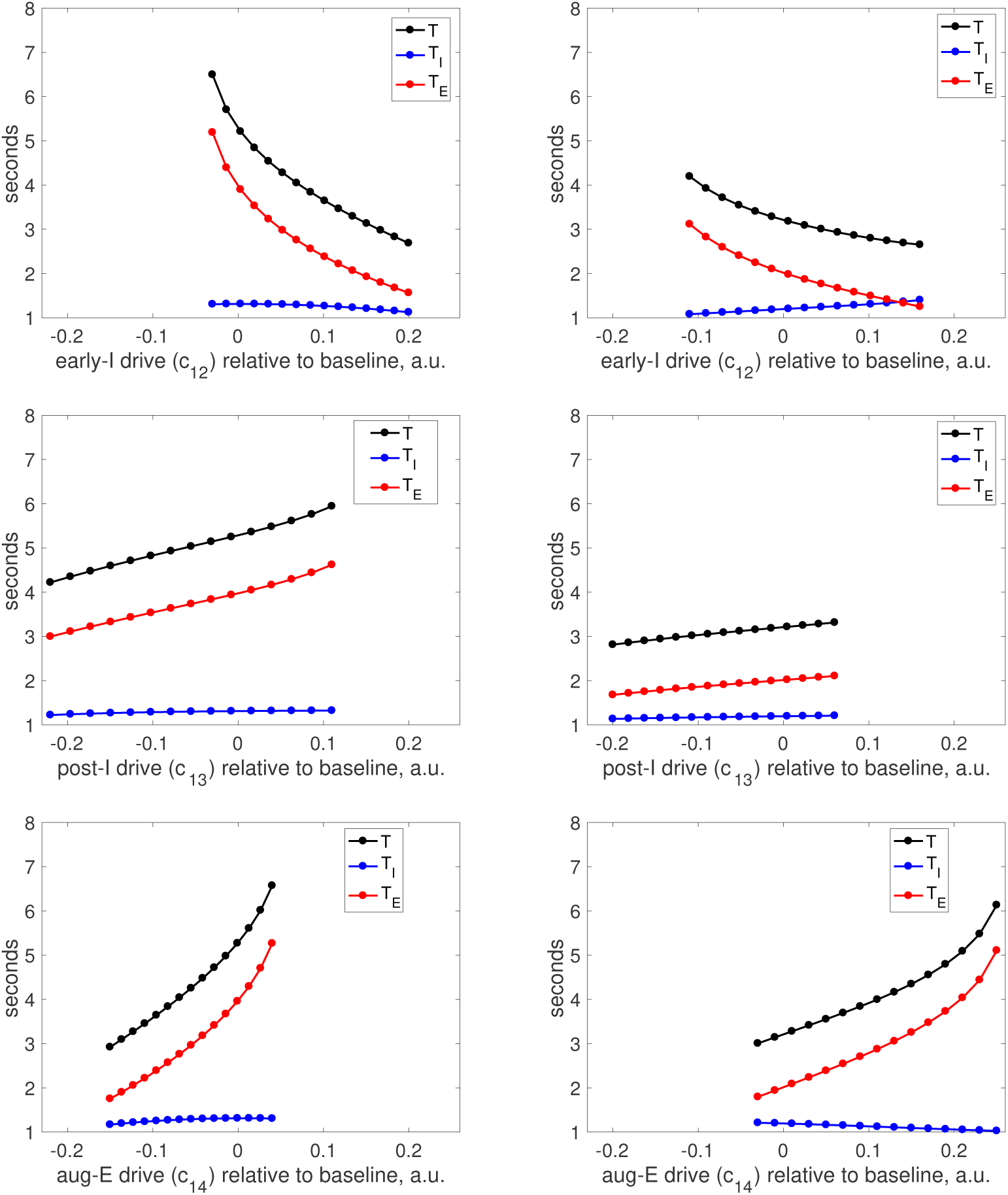
Inputs to different nodes in the network provide distinct period control mechanisms. Left column: pre-I is tuned to be intrinsically oscillatory (*c*_11_ = *−*0.03). Right column: pre-I is tuned to be intrinsically tonic (*c*_11_ = 0.01). All panels show period (black curve), inspiratory duration (blue curve), and expiratory duration (red curve) versus the value of a drive parameter relative to its baseline value (here labeled as 0) that was varied over the full range that preserved network rhythmicity. Drive parameters used are drive to early-I, *c*_12_ (top), drive to post-I, *c*_13_ (middle), and drive to aug-E, *c*_14_ (bottom). Drive parameters are fixed at *c*_12_ = 0.19, *c*_13_ = 0.58, *c*_14_ = 0.2 unless being varied. The same horizontal and vertical axis ranges are used in all panels.

### 2.5 Network responses to changes in synaptic inhibition

Recent experiments considered perturbations of respiratory rhythms resulting from microinjection of GABAA receptor and glycinergic receptor antagonists into the preBötC or the BötC of rats to disrupt post-synaptic inhibition [24]. The former manipulation led to a decrease in respiratory period and amplitude, as measured by integrated phrenic nerve activity. In an *in situ* rat brainstem-spinal cord perfused preparation, sufficient blockade of inhibition to the preBötC completely disrupted the respiratory rhythm. Sufficient blockade of inhibition to the BötC, where the subpopulations of post-I and aug-E inhibitory neurons providing input to preBötC circuits are thought to reside [8, 15, 24], also perturbed respiratory rhythms, in both *in vivo* and *in situ* experiments. In contrast to the experiments on the preBötC, however, this manipulation increased respiratory period while decreasing the amplitude of phrenic nerve outputs. In some cases, rhythmic activity was completely suppressed with block of inhibition in the BötC. Responses of our model to reductions in inhibition were similar, but not identical, to the experimental findings.

Our results agreed completely with the experimental findings [24] associated with reduction of inhibition to the pre-I and early-I units of the preBötC, both for intrinsically oscillatory and for intrinsically tonically active pre-I tunings. Indeed, in both cases, period was approximately halved and amplitude, measured in terms of inspiratory output, dropped by about 40% (Fig. 10A,B), quantitatively consistent with the experimental results. As observed *in vivo*, the drop in period included shortening of both *T*_*I*_ and, especially, *T*_*E*_. The reduction in *T*_*E*_ was clearly expected based on the lessened ability of the expiratory units to suppress the inspiratory units, while the decrease of *T*_*I*_ resulted from the fact that inspiratory onset occurred with less deinactivation of *I*_*NaP*_, yielding a shorter time needed for *I*_*NaP*_ inactivation to occur. The production of a rhythmic output failed at a higher remaining fraction of inhibition when the pre-I unit was intrinsically tonic than when it was oscillatory, because the pre-I and early-I units became tonically active within the network with less of a loss of inhibition than in the former case.

**Fig 10.**
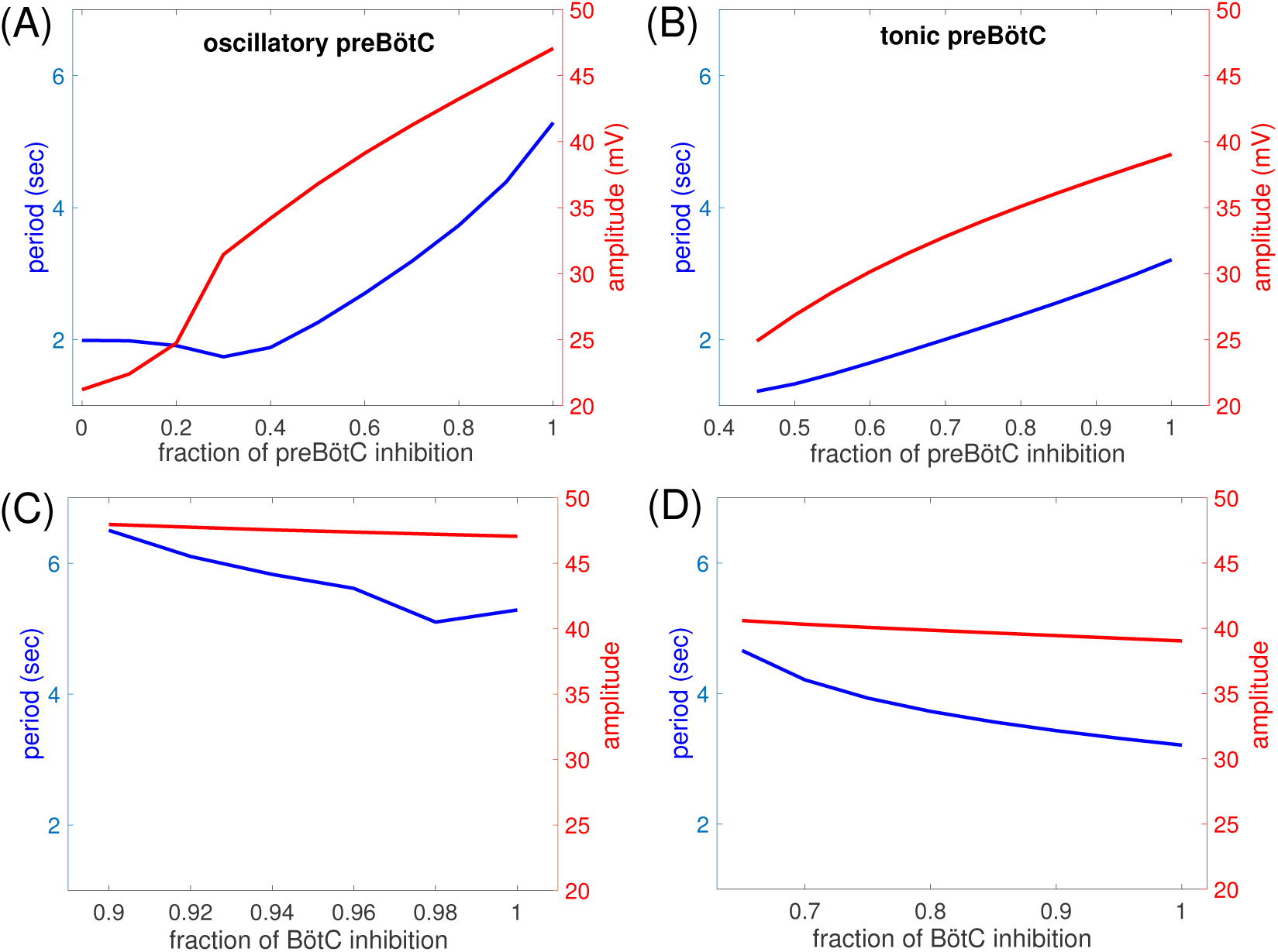
Dependence of respiratory period and amplitude of inspiratory signal on inhibition levels. (A) pre-I unit is intrinsically oscillatory (*c*_11_ = –0.03) and inhibition to the preBötC units (pre-I and early-I) is varied. (B) pre-I unit is intrinsically tonic (*c*_11_ = 0.01) and inhibition to the preBötC units (pre-I and early-I) is varied. (C) pre-I unit is intrinsically oscillatory (*c*_11_ = *−*0.03) and inhibition to the BötC units (post-I and aug-E) is varied. (D) pre-I unit is intrinsically tonic (*c*_11_ = 0.01) and inhibition to the BötC units (post-I and aug-E) is varied. All horizontal axis ranges are distinct; in all cases, the baseline network corresponds to 1 and the lower bound denotes a level just below which the oscillation is lost. The network response to lowered inhibition is qualitatively similar regardless of whether the pre-I unit is oscillatory or tonic. Period and amplitude are both much more strongly modulated by changes in inhibition to the preBötC units than by changes in inhibition to the BötC units; note also that the slopes of the period curves are opposite in the two cases.

When we decreased inhibition to the post-I and aug-E units, corresponding to experimental perturbations of synaptic inhibition in the BötC, the network did not exhibit an appreciable change in inspiratory output amplitude. When the pre-I unit was intrinsically oscillatory, output period was relatively insensitive to the inhibition level to the BötC within the range over which rhythmicity persisted, although it did increase by about 20% (Fig. 10C). With a reduction of inhibition of only about 10%, output rhythmicity was lost. An intrinsically tonic tuning of the pre-I unit provided more robustness to the stronger inhibition from expiratory units that resulted from a decrease in inhibition to the BötC, such that network rhythmicity persisted down to about 65% of normal inhibition levels. Moreover, network period increased by almost 50% over this range of inhibition levels, due to the same effect that was observed to be dominant experimentally, namely an increase in *T*_*E*_, consistent with the mechanism of network rhythmogenesis (Fig. 10D). This comparison with experimental results supports the idea that an intrinsically tonic tuning of pre-I neurons may be more consistent with experimental observations than an intrinsically pacemaking tuning, since the tonic tuning provides more robustness against decreased inhibition in the BötC [24].

### 2.6 Network rhythmicity remains similar when an excitatory post-inspiratory complex (PiCo) unit is included

Recently, a glutamatergic post-inspiratory complex (PiCo) including neurons with intrinsic oscillatory capabilities has been identified [16]. As its name suggests, PiCo neurons activate during the post-inspiratory phase. The timing of PiCo activation shifts to inspiration when the inhibition that it normally receives during the I phase is blocked, suggesting that the PiCo receives excitatory inputs from glutamatergic inspiratory neurons. To explore its possible role, we incorporated an intrinsically oscillatory PiCo unit into our model network, sending excitation to the post-I unit, excited by the pre-I unit, and, like the post-I inhibitory unit, inhibited by the early-I and aug-E units. We modeled the PiCo as an intrinsically oscillatory unit [16], with a voltage equation including the same formulation of *I*_*NaP*_ used in the pre-I unit and, for simplicity, with the same persistent sodium and leak conductances as the pre-I unit as well.

The inclusion of the PiCo did not qualitatively alter network activity patterns: the PiCo remained suppressed by inhibition from early-I during inspiration and activated together with post-I during the post-inspiratory phase (Fig. 4C,D). Excitatory output from the PiCo promoted post-I activation, causing the post-I unit to exhibit less adaptation during its active phase and, correspondingly, making the activation of the aug-E unit slightly less gradual (compare Figure 4C,D vs. Figure 4A,B). Because the network was already rhythmic without the presence of the PiCo, turning on and strengthening the excitation from the PiCo to the post-I unit had little effect on expiratory phase duration, regardless of whether the pre-I unit was intrinsically oscillatory or tonic (Fig. 11A,B). Examining this effect in more detail, we observe that excitation from PiCo to the post-I unit induces a stronger post-I inhibition of its postsynaptic targets, and of pre-I in particular, while PiCo is active. As a result of this stronger inhibition, the persistent sodium current for the pre-I unit deinactivates more than it would otherwise during the expiratory phase. After the post-I unit falls silent, the pre-I is left with approximately the same inhibitory input (from the aug-E unit) as it would have been without the PiCo, but with slightly larger *h*_*pre–I*_ (Fig. 11C,D; compare grey and purple circles). With similar inhibition levels, a similar level of persistent sodium deinactivation is needed to allow pre-I activation; with PiCo, since the value of *h*_*pre–I*_ at the start of aug-E is larger, the augmenting-expiratory phase ends up being slightly shorter, resulting in a mildly shorter overall cycle period.

**Fig 11.**
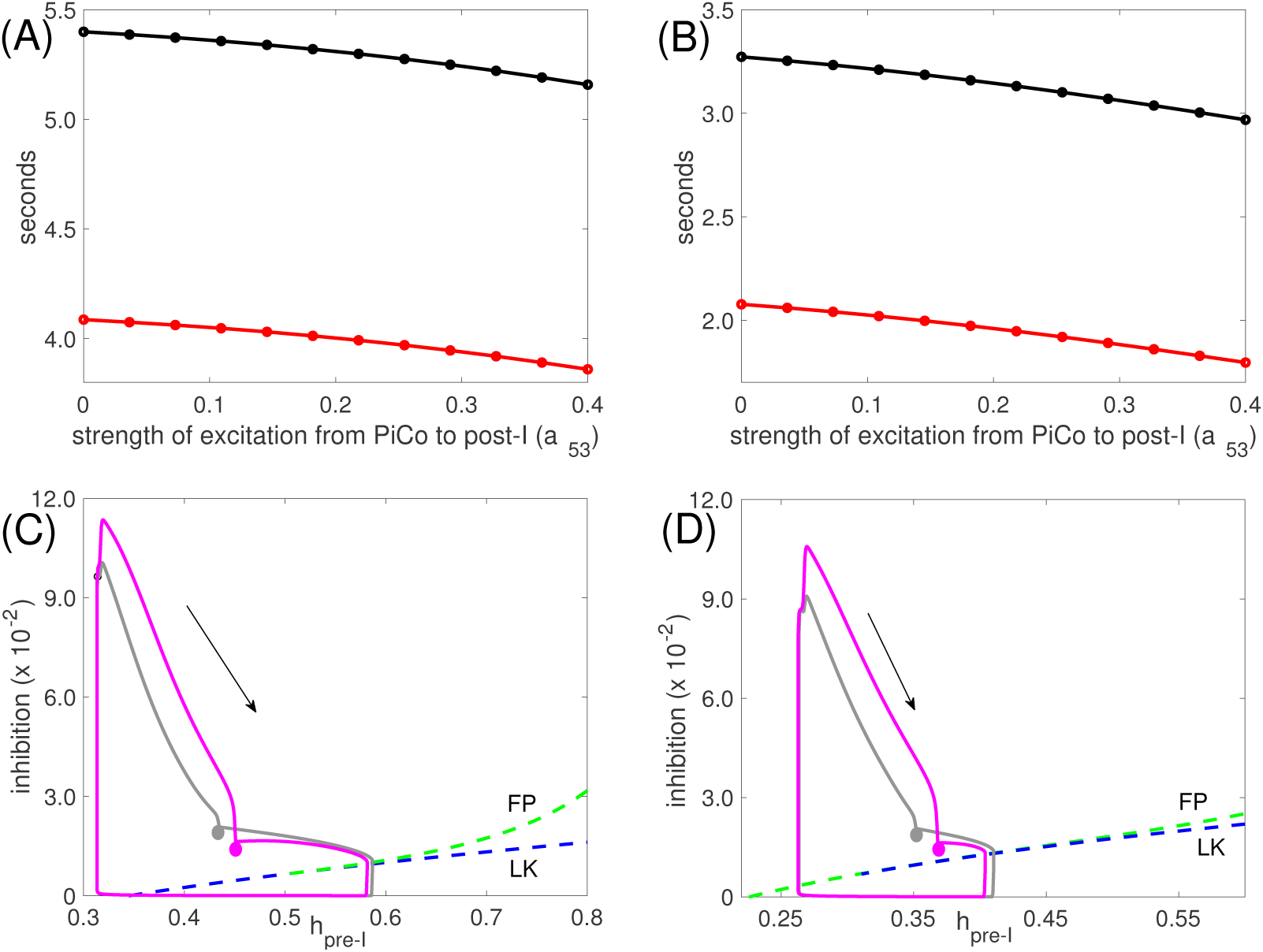
Effect of increasing drive from PiCo to the post-I unit (*a*_53_) on respiratory period. (A) Period (black) and expiratory phase duration (red) for *c*_11_ =–0.03 (pre-I unit intrinsically oscillatory). (B) Period (black) and expiratory phase duration (red) for *c*_11_ = 0.01 (pre-I tonic). Note that the entire change in period is captured by the change in expiratory phase duration in both (A) and (B). (C) Trajectories projected into the slow phase plane for the pre-I unit with *c*_11_ = *−*0.03 for *a*_53_ = 0 (without PiCo activity, grey) and *a*_53_ = 0.4 (with PiCo activity, purple). The curves of fixed points (green, FP) and left knees (blue, LK) for the pre-I unit dynamics are also shown. (D) Trajectories projected into the slow phase plane for the pre-I unit with *c*_11_ = 0.01 for *a*_53_ = 0 (grey) and *a*_53_ = 0.4 (purple). FP and LK as in (C). In both (C) and (D), the projected trajectories reach the points marked with circles at the start of the aug-E phase. The arrow indicates the direction of flow.

Given that the PiCo unit oscillates and drives post-I, it seems theoretically possible that if the pre-I unit is tuned to produce tonic inspiratory activity, the PiCo could conceivably rescue the network rhythm by activating and recruiting post-I, which could in turn shut down the pre-I and early-I units. In our network configuration, however, sustained pre-I activity supports sustained early-I activity, which suppresses PiCo, such that the rescue fails. If inhibition from early-I to PiCo is weakened, then depending on how parameters are tuned, either PiCo joins pre-I/early-I in sustained activation, PiCo oscillates without recruiting post-I and hence without interrupting inspiratory activity, or other pathological patterns emerge. On the other hand, if post-I is tuned to be tonically active and to suppress inspiration without PiCo, then rhythmic drive from PiCo to post-I does not terminate post-I activity to rescue inspiration. Extensive parameter exploration did not allow PiCo to rescue network oscillations; if the PiCo is able to activate in the regime where the network is non-oscillatory without it, then it fails to maintain its post-I timing or restore rhythmic network activity patterns. Thus, although this issue needs to be explored further, our findings support the conjecture that a post-inspiratory oscillatory unit cannot on its own induce network oscillations in an otherwise non-rhythmic respiratory network.

Finally, we examined how tuning the drive to the PiCo unit (*c*_15_) impacted network outputs and how the presence of the PiCo affected network responses to tuning of drives to other units (*c*_1*i*_, *i <* 5). Varying *c*_15_ switched intrinsic PiCo dynamics from complete quiescence (stable fixed point at hyperpolarized voltage) to oscillations to tonic activation (stable fixed point at depolarized voltage; Fig. 12A,B), as is typical for the voltage-dependent *I*_*NaP*_ -mediated oscillation mechanism. For small enough *c*_15_, even excitation from pre-I could not induce PiCo recruitment at or near the onset of the post-I phase on all rhythm cycles, while for large enough *c*_15_, PiCo excitatory output allows post-I to suppress aug-E throughout expiration. Between these extremes, *c*_15_ had little effect on network period (Fig. 12A,B). Although variation of *c*_15_ altered the projection of the network output trajectory to the PiCo phase plane, the PiCo active phase duration remained essentially invariant (data not shown). Larger *c*_15_ produced slightly stronger PiCo output and hence affected post-I output quantitatively, but PiCo deactivation remained associated with a signficant drop in post-I output and corresponding emergence of aug-E activity, with little change in overall timing (∼ 200 msec change over the full range of *c*_15_). Similarly, because PiCo activation remained stereotyped, its inclusion produced little impact on network responses to other drive parameters (cf. Fig. 12C,D, and compare to Fig. 9, middle row). One subtle effect that emerged from inclusion of the PiCo is that its output could support post-I activation for lower levels of *c*_13_ than otherwise possible, and, interestingly, could preserve rhythmic network outputs for slightly higher levels of *c*_13_ as well (Fig. 12C,D and Fig. 9, middle row, compare range of *c*_13_ values), through effects of inhibition on *I*_*NaP*_.

**Fig 12.**
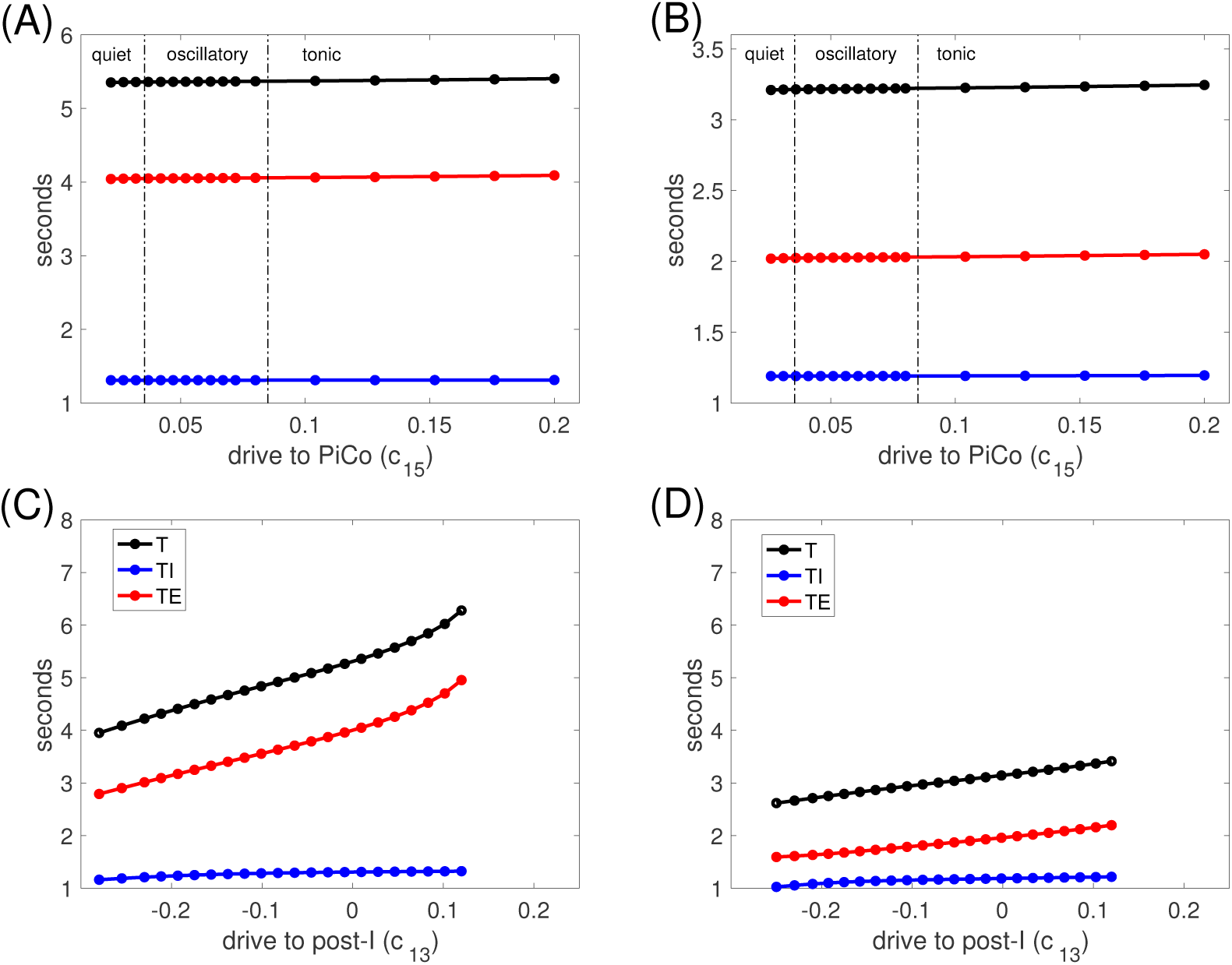
Period is insensitive to drive to the PiCo and to the presence of PiCo. Period *T* (black), expiratory duration *T*_*E*_ (red), and inspiratory duration *T*_*I*_ (blue) versus (A,B) PiCo drive parameter (*c*_15_) or (C,D) drive to the post-I unit (*c*_13_), relative to its baseline value of 0.58 (denoted by 0 here). In (A,B), dashed-dotted vertical black lines separate intervals of *c*_15_ values corresponding to different types of PiCo intrinsic dynamics. In (A,C), pre-I is intrinsically oscillatory (*c*_11_ = *−*0.03); in (B,D), pre-I is intrinsically tonic (*c*_11_ = 0.01). Axis ranges in (C,D) are identical to those in Fig. 9 (middle row) except for extension to more negative relative values of *c*_13_ here, since rhythmicity persists down to these lower levels.

## 3 Discussion

In this work, we considered a computational network of neuronal units connected synaptically and receiving tonic excitatory synaptic drive, with network parameters tuned to produce a multi-phase rhythmic activity pattern, as a model of components of the mammalian respiratory CPG. Our most fundamental result was the observation that network outputs persist across regimes of tuning of a conditionally rhythmic unit in the network, representing the inspiratory pre-I oscillator in the preBötC. This robustness and insensitivity of qualitative network output pattern to pre-I intrinsic dynamics was not inevitable but also did not require special tuning. These properties emerged because synaptic interactions shaped the network dynamics, altering the pre-I unit’s outputs compared to what they would be in the isolated preBötC and resulting in similar network dynamic mechanisms across intrinsic tuning regimes for pre-I. While qualitatively similar network outputs were maintained, the ability of the network to function across a broad range of pre-I excitability resulted in an extended dynamic range of output amplitude and period, which could be valuable for function across diverse conditions.

Our findings can be viewed as exposing general properties of networks incorporating conditional oscillators under certain conditions and are also of specific relevance to an ongoing debate about rhythmicity in the respiratory CPG, which motivated our choices of network connectivity and unit dynamics to study. In the respiratory context, our results point out that a CPG network structure that has been hypothesized based on experimental data [5, 10] yields network dynamics that are robust across a range of tunings of the key glutamatergic respiratory neurons in the preBötC. Our findings also imply that experimental manipulations that would alter dynamics or excitability in the isolated preBötC [4, 6, 21, 25, 26] cannot be expected to yield analogous results when applied to the connected respiratory network, and that the failure of such manipulations to compromise respiratory outputs does not imply a lack of a role for pre-I dynamics in respiratory rhythm generation. Moreover, our modeling analyses extend our previous results [10] indicating that all of the units participating in rhythmic respiratory activity that are proposed to be core elements of the respiratory CPG can represent important nodes for the control of rhythm features. This control appears to occur seamlessly without altering the basic three-phase organization of activity within the CPG, consistent with experimental observations [15, 26] and the notion that input tuning of excitability of different circuit elements is robust in terms of maintaining the coordinated activity necessary for proper behavioral function.

More specifically, our work predicts that increases in tonic drive to pre-I and early-I respiratory neurons should shorten the respiratory cycle period via decreases in the duration of expiration, which is consistent with experimental results [15, 21, 26]. Increases in tonic drive to post-I and aug-E respiratory neurons are predicted to increase the period, again via changes in expiratory duration, also consistent with experimental observations [15]. We found that tuning of the aug-E population produces a broader range of periods than tuning of post-I; aug-E output in baseline conditions is far from maximal levels, due to lingering inhibition from post-I, and hence increased drive to aug-E can have a significant impact on the timing of inspiratory onset. We also predict that the maximal amplitude of pre-I outputs will be larger in the connected network than in the isolated preBötC, while the minimal network amplitude will also exceed the minimal preBötC amplitude (Fig. 2). The inclusion of *I*_*NaP*_ in our pre-I model yields oscillations with small amplitude and short period when pre-I excitability is high, as well as very long oscillations just before the transition to quiescence as excitability drops, in the isolated pre-I population. While the connected network cannot capture this full range of periods, the fact that it continues to produce functional outputs across different regimes of pre-I intrinsic dynamics significantly extends its dynamic range relative to what it could produce if restricted to a specific pre-I tuning. Interestingly, our model produces some evidence in support of the idea that in conditions associated with respiratory outputs in the *in situ* perfused rat preparation, pre-I respiratory neurons are tuned to be tonically active, based on model responses to decreases in *g*_*NaP*_ used to simulate partial *I*_*NaP*_ blockade (Fig. 5). We also found that the tonic pre-I tuning gives more robustness of rhythmic network activity to decreases in inhibition to BötC neurons than does the oscillatory tuning (Fig. 10), because the tonic tuning allows inspiratory neurons to activate despite receiving stronger inhibitory outputs from the BötC. Note, however, that a tonic pre-I tuning does not imply that *I*_*NaP*_ does not play any role in pre-I activity nor that it is completely inactivated. On the other hand, we observed that the oscillatory pre-I tuning generally could yield longer respiratory cycle periods (hence extended dynamic range) and associated expiratory phase durations over changes in various drive and synaptic parameters (Fig. 9), because with this tuning, the escape of the pre-I unit from inhibition, needed to initiate inspiration, could become strongly delayed. The oscillatory tuning also gave more robustness to decreases in inhibition to preBötC neurons because this tuning yielded more protection against tonic activation of pre-I and early-I within the full network (Fig. 10). It is therefore possible that neuromodulatory adjustment of pre-I excitability could be used to transition the network between these regimes, each allowing for particular network tuning mechanisms to become accessible.

Related results to ours have recently been demonstrated in a half-center oscillator network, using a similar reduced model that was shown to capture many properties found in simulations of a network composed of a large collection of spiking Hodgkin-Huxley style model neurons [27]. A key point in that work was that rather than the intrinsic dynamics of individual units in the network, the dynamic transition mechanisms by which switches in active populations occurred, involving synaptic effects known as escape and release [10, 28-30], determined network output properties. This finding is consistent with our proposal that masking of pre-I properties arises because the dynamic regimes set up by network interactions are similar across pre-I unit tunings.

The ideas that we explore are consistent with arguments made by Richter and Smith [8] that in the intact respiratory circuit, endogenous pacemaker currents of preBötC neurons need not be autonomously active to drive respiration and indeed may not be active during rhythm generation by the full respiratory CPG circuit due to control of membrane potentials by inhibitory synaptic currents from network connections from the BötC; furthermore, the frequency of the full rhythm may not match the intrinsic frequency of pacemaker neurons in isolation. Diekman et al. have also considered the related ideas that distinct mechanisms may underlie network bursting under different feedback conditions and that an intrinsic tuning of preBötC neurons that does not favor bursting may enhance the robustness of network bursting across normoxic and hypoxic regimes in the presence of chemosensation [31].

Our results are also in the same spirit as many years of experimental and modeling results from the CPG in the crustacean stomatogastric ganglion (STG), which generates digestive rhythms. These experiments and simulations have shown that network patterning in the STG is similar across wide ranges of values of parameters associated with the neurons in the circuit [32, 33]. Our work carries this idea to the respiratory network and explores additional issues related to the two forms of masking that we have discussed. In particular, we demonstrate that changes in parameter values that alter the intrinsic dynamics of a critical node in a CPG if applied in isolation, for example by inducing a bifurcation, need not produce any noticeable impact on the dynamics of a network in which that node is embedded. Moreover, once that node is embedded in the network, the influence of the network can cause its output tuning to change, effectively masking or overriding its intrinsic dynamics due to circuit interactions.

We performed some simulations with the newly discovered oscillatory post-I population, the PiCo [16], included in our model network, receiving excitatory input from pre-I and inhibition from early-I and aug-E, while providing excitation to the post-I inhibitory unit, based on the proposal that PiCo is critical for generation of post-I activity [16]. Interestingly, our simulations predict that changes in drive to the PiCo have no effect on respiratory period and increases in the strength of excitatory drive from the PiCo to post-I only slightly alter the period, causing it to decrease by a small amount (Fig. 11). We modeled the intrinsic dynamics of the PiCo identically to the dynamics of the pre-I unit, based on observations of its intrinsic rhythmicity [16], although it remains for future experiments to test the mechanisms underlying this rhythmicity. Our results raise the question: in what way does the presence of PiCo enhance respiratory network function? According to our simulations, PiCo has limited effectiveness in restoring network rhythmicity if rhythmicity is not present without PiCo, but perhaps PiCo could serve a rescue function in some other circumstances that we have not explored. For example, Anderson et al. did demonstrate an impact on network dynamics by showing that PiCo photostimulation leads to vagal bursts and delays subsequent inspiration [16]. These results are consistent with the idea that stimulating PiCo excitatory neurons drives post-I motor output and can also drive post-I inhibitory neurons to effect the dynamic transition from expiration to inspiration as proposed in our model. The recently proposed triple oscillator hypothesis includes PiCo as a critical component, linked via excitatory coupling with preBötC rhythm-generating neurons and the parafacial respiratory group (pFRG) conditional oscillator, which generates late expiratory activity [34]. We have not included in the present analysis the pFRG oscillator, which is not active and hence not involved in generating the aug-E unit activity during normal eupneic breathing. We have previously (e.g., [35]) analyzed in detail network dynamics when the pFRG and pre-I intrinsic oscillators are coupled and their activity is coordinated by a more extensive inhibitory connectome than represented by the present model, which is intended to represent generation of the normal three-phase rhythmic pattern during eupneic breathing. An interesting future direction would be to analyze the network dynamics in an extended model with coupling between pre-I, PiCo, and pFRG intrinsic oscillators and to evaluate how these interactions impact network performance in conditions (e.g., elevated CO_2_) under which the pFRG is functionally active.

Our results were based on simulations of a network of units with two-dimensional dynamics, connected via synaptic coupling. This framework is perfectly valid for illustrating the fundamental point that intrinsic dynamics can become masked by network interactions. It makes the link with the respiratory CPG more abstract, however. To make this connection, we think of each unit within the network as representing the average activity of a population with synchronized activation and inactivation but with asynchronous spiking. The assumed synchrony within the preBötC is consistent with observed eupneic dynamics, presumably reflecting a lack of strong, destabilizing inhibition intrinsic to the preBötC [36]. The model does include the key features of conditional autorhythmicity of the excitatory unit, along with adaptation of the inhibitory units. Furthermore, past work has established that it can produce similar output patterns and even similar responses to changes in drives and coupling to a much more detailed model involving many coupled Hodgkin-Huxley neurons [10, 27, 37]. Agreement of pre-I outputs with experimental recordings of inspiratory (pre)motor nerve outputs also supports the use of this framework.

We based the conditional intrinsic rhythmicity in our model pre-I unit on the persistent sodium current, *I*_*NaP*_, a well studied current known to exist and to support the possibility of rhythmogenesis in preBötC excitatory neurons. Our results immediately generalize, however, if the pre-I unit instead features other currents or combinations of currents that include one amplifying factor and one resonant (or depolarization-resisting) factor acting on appropriate timescales [14], such that increasing drive yields transitions from quiescence to oscillations to sustained activation via A-H bifurcations (Fig. 2B). They may also apply to scenarios where individual pre-I neurons cannot become rhythmic but instead the collective pre-I population can be tuned to produce rhythmicity, either through expansion of dynamic range due to synaptic coupling (e.g., [38-40]) or through inclusion of various ionic currents (e.g., [8, 41, 42]), if the bifurcation structure associated with the activity transitions is similar to what we have considered. Nonetheless, specific predictions about the respiratory network, especially at the quantitative level, may suffer from the abstraction in the model, as well as from the omission of other currents, including the CAN current, which have been shown to contribute to respiratory outputs [43-45]. Our model also cannot represent the heterogeneity that is naturally present within biological populations of neurons, which has been predicted to add robustness to respiratory outputs [38-40]. Indeed, with heterogeneity, it is likely that different neurons have different ionic conductances leading to different intrinsic dynamics [40]. In this case, a generalization of our conclusions would be that the precise distribution of these intrinsic dynamic properties is not critical to the characteristics of the full network outputs. Another caveat of our results is that we saw loss of network rhythmicity with sufficient block of inhibition in all regimes considered, whereas outcomes from experiments involving inhibitory block have been more diverse [22, 24, 46]. In reality, experiments can show variable perturbations from disruption of inhibition (e.g., whether or not rhythm is disrupted) across different experimental trials, possibly corresponding to different levels of inhibition block achieved in different experiments.

### 3.1 Conclusion

CPGs subserve many functions critical to survival, including mammalian respiration. It is critical for functional outputs to persist across wide ranges of metabolic and environmental conditions including diverse developmental and disease states. Maintenance of function as well as control of outputs to allow effective behavior across conditions likely involve significant alterations in feedback and top-down drives to network components and in neuromodulatory influences [47]. Hence, even if particular components within a CPG circuit exhibit intrinsic rhythmicity for certain drive and neuromodulator levels, their dynamics may change as these levels vary. Therefore, the invariance of network outputs and tuning under changes in intrinsic dynamics that we highlight is likely a critical feature of maintaining stable CPG performance across conditions. In the case of the respiratory network, this invariance ensures flexible and robust performance over a wide dynamic range of rhythm generation, which is likely a critical property for such a vital homeostatic CPG.

## 4 Methods

All results in this paper were derived from simulations of highly reduced ordinary differential equation models for the excitatory kernel of the preBötC and for a respiratory CPG network that includes this preBötC component as well as inhibitory early-I, post-I, and aug-E components and, in some simulations, an additional excitatory PiCo component (Fig. 1). The models were based on those used in several previous respiratory modeling studies [10, 35, 37, 48]. In this framework, each neuronal population is assumed to undergo synchronized transitions between active and silent states, but without spike-level synchrony. Thus, each is represented by a single non-spiking neuron model unit, the voltage of which corresponds to the average of the membrane potentials of the neurons in that population.

For purposes of model specification, we number the model units as follows: 1 - pre-I, 2 - early-I, 3 - post-I, 4 - aug-E, 5 - PiCo. All units obeyed the equations

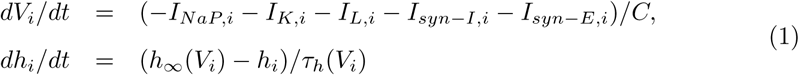

where *i* ∈ {1,*…,* 5}. In system (1), *I*_*NaP,i*_ represents the current through persistent sodium channels, *I*_*K,i*_ is a potassium current, *I*_*L,i*_ = *g*_*L,α*_*·* (*V*_*i*_ *- E*_*L,α*_) is the leakage current with *α* = *exc* for *i* ∈ {1, 5} and *α* = *inh* for *i* ∈ {2, 3, 4}, respectively. Moreover, *I*_*syn-I,i*_ and *I*_*syn-E,i*_ are inhibitory and excitatory synaptic currents, respectively.

For the excitatory units, with *i* ∈ {1, 5}, the intrinsic currents in (1) take the following forms:

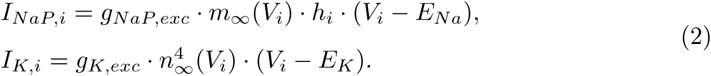

For the inhibitory units, with *i* ∈ {2, 3, 4}, the intrinsic currents in (1) are given by:

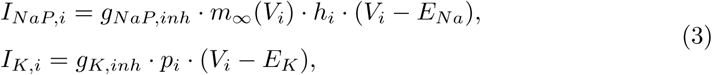

with

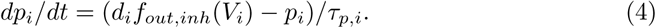

That is, the persistent sodium current takes the same form for all units but has a stronger conductance for excitatory than inhibitory units. On the other hand, as in previous models (e.g., [10]), the potassium current for the excitatory units is a standard delayed rectifier that is set to be quite weak, representing potassium current activated by the depolarization attributable to the persistent sodium current, while for the inhibitory units it is a much stronger adaptation current with a dedicated gating variable, *p*_*i*_. The voltage-dependent activation functions and time constants in equations (1), (2), (3) are described by the following functions:

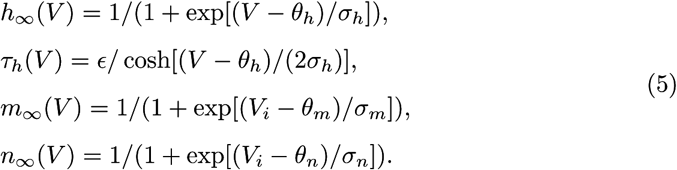

In equation (4) for the adaptation current gating variable, the function *f*_*out,inh*_(*V*) appears. This function represents a nonlinear filtering of the output of a unit and takes the sigmoidal form

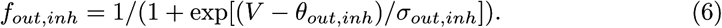

Thus, equation (4) specifies how the conductance of an inhibitory unit’s adaptation current grows while that unit is active.

To define the synaptic currents in equation (1), we use the function given in equation (6) along with a similar sigmoidal function

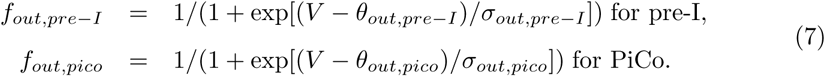

The synaptic currents to unit *i* are given by

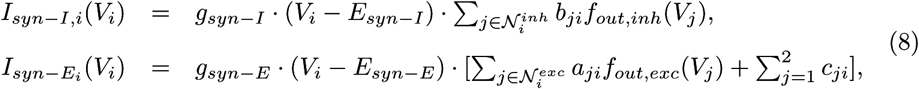

where 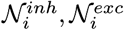 denote the sets of inhibitory and excitatory units, respectively, presynaptic to unit *i*, the constants *b*_*ji*_, *a*_*ji*_ give the strengths of inhibitory and excitatory connections, and the terms *c*_*ji*_, *j* ∈ {1, 2}, are parameters representing the strength of the tonic drive to each unit, with *c*_1*i*_ tunable and *c*_2*i*_ fixed for each *i*.

A baseline parameter set was used for all results unless otherwise specified in the text. This set included the following values:

- **pre-I and PiCo:** *g*_*NaP,exc*_ = 4.5*nS, g*_*K,exc*_ = 1.0*nS, g*_*L,exc*_ = 3*nS, E*_*L,exc*_ = *−*65*mV, θ*_*out,pre-I*_ = *−*32*mV, σ*_*out,pre-I*_ = *−*8*mV, θ*_*out,pico*_ = *−*20*mV, σ*_*out,pico*_ = *−*12*mV*
- **early-I, post-I, and aug-E:** *g*_*NaP,inh*_ = 0.25*nS, g*_*K,inh*_ = 10.0*nS, g*_*L,inh*_ = 3.25*nS, E*_*L,inh*_ = *−*60*mV, θ*_*out,inh*_ = *−*30*mV, σ*_*out,inh*_ = *−*4*mV, τ*_*p,*2_ = 2000*ms* (early-I), *τ*_*p,*3_ = 1500*ms* (post-I), *τ*_*p,*4_ = 2000*ms* (aug-E)
- **shared:** *C* = 20*pF, E*_*Na*_ = 50*mV, E*_*K*_ = *−*85*mV, θ*_*h*_ = *−*48*mV, σ*_*h*_ = 8*mV, E* = 4000*ms, θ*_*m*_ = *−*37*mV, σ*_*m*_ = *−*6*mV, θ*_*n*_ = *−*29*mV, σ*_*n*_ = *−*4*mV, g*_*syn-E*_ = 10*nS, g*_*syn-I*_ = 60*nS, E*_*syn-E*_ = 0*mV, E*_*syn-I*_ = *−*75*mV*

Additional synaptic connection strength parameter values for each unit as specified in equation (8) were as follows:

- **pre-I (unit 1):** *b*_3,1_ = 0.125, *b*_4,1_ = 0.015, *c*_2,1_ = 0.095, with *c*_1,1_ values given in the text
- **early-I (unit 2):** *a*_1,2_ = 0.6, *b*_3,2_ = 0.27, *b*_4,2_ = 0.3, *c*_1,2_ = 0.19, *c*_2,2_ = 0.3
- **post-I (unit 3):** *a*_5,3_ = 0.1 (when PiCo was included), *b*_2,3_ = 0.6, *b*_4,3_ = 0.05 without PiCo or 0.02 with PiCo, *c*_1,3_ = 0.58, *c*_2,3_ = 0
- **aug-E (unit 4):** *b*_2,4_ = 0.3, *b*_3,4_ = 0.45, *c*_1,4_ = 0.2, *c*_2,4_ = 0.4
- **PiCo (unit 5):** *a*_1,5_ = 0.2, *b*_2,5_ = 0.2, *b*_4,5_ = 0.3, *c*_1,5_ = 0.045

All simulations were run in XPPAUT [20] and Matlab (The Mathworks, Inc., Natick, MA, USA) and the latter was used to generate computational figures. Based on these simulations, we define the parameter ranges over which the network generates functional rhythmic outputs with the correct order of activation (pre-I activation before or together with early-I activation, followed by post-I activation and then aug-E activation in each cycle) and with greater aug-E output than post-I output at the end of the expiratory phase (*f*_*out,inh*_(*V*_4_) *> f*_*out,inh*_(*V*_3_); e.g,. Fig. 4).

Some figures in this paper include nullclines or bifurcation diagrams. Note that system (1) for a single unit in the excitatory case (i.e., without a *p* variable) consists of a pair of coupled differential equations with dependent variables *V, h*, subject to time-dependent inputs from the synaptic currents *I*_*syn-I*_, *I*_*syn-E*_. If these inputs are held fixed or are set to zero to represent a unit in isolation, then the behavior of solutions of the system can be visualized in the (*V, h*) plane. Within this plane, the set of (*V, h*) for which *dV/dt* = 0 holds is the *V* -nullcline, with an analogous definition for the *h*-nullcline. For system (1), each of these sets is a curve (Fig. 2A). In particular, for a certain range of parameters, the *V* -nullcline is a cubic-shaped curve with two folds, or knees, the left knee (LK) and right knee (RK).

The position of the each nullcline will vary with changes in model parameters that appear in the corresponding ODE. Intersection points of the nullclines are equilibria of the system; if (*V, h*) are initially chosen to correspond to their values at an equilibrium point, then (*V, h*) will remain constant for all time. For sufficiently small *c*_11_, the equilibrium point (or fixed point, FP) lies on the left branch of the *V* -nullcline. Yet since the *V* -nullcline for the pre-I unit moves under modulation of *c*_11_ (Fig. 2A), the position of the unit’s equilibrium point also changes. We use XPPAUT to generate a bifurcation diagram for the pre-I unit by tracking both the location and the stability of the equilibrium point (Fig. 2B), along with information about periodic orbits that emerge (disappear) when the equilibrium loses (gains) stability, as *c*_11_ is increased.

As *c*_11_ is increased, the *c*_11_ value where the pre-I equilibrium loses stability represents the transition from quiescent to oscillatory intrinsic dynamics for our baseline parameter set, while the *c*_11_ value where the equilibrium regains stability corresponds to the transition from oscillatory to tonic intrinsic dynamics. If any other parameter from the pre-I system is altered, then this variation may change these transitional *c*_11_ values, Thus, for each transition, we can use XPPAUT to obtain a curve of values in the two-dimensional (*c*_11_, *g*_*NaP,exc*_) parameter space where this transition occurs; the curve corresponding to the oscillatory-to-tonic transition appears in each panel of Fig. 5.

In the description above, we set the inhibition level to the pre-I unit equal to 0. We can also treat that inhibition level as a parameter of the pre-I model. Variation in the inhibition level will alter the position of the *V* -nullcline and hence of the LK and FP. We also use XPPAUT to compute how these features depend on the inhibition level, with results appearing in Figs. 6, 7, 11.

## Acknowledgments

JR acknowledges support from NSF awards DMS 1312508 amd DMS 1612913. This work was also supported in part by the Intramural Research Program of the NIH, NINDS. The authors thank Mohammad Tariq for assistance and discussions in the early stages of this project.

